# Reimplementation of the Potjans-Diesmann cortical microcircuit model: from NEST to Brian

**DOI:** 10.1101/248401

**Authors:** Renan Oliveira Shimoura, Nilton Liuji Kamiji, Rodrigo Felipe de Oliveira Pena, Vinicius Lima Cordeiro, Cesar Celis Ceballos, Cecilia Romaro, Antonio Carlos Roque

## Abstract

This work targets the replicability of computational models to provide the community with tested and proven open-source models to be used in new studies and implementations. The Potjans-Diesmann model describes a cortical microcircuit containing two cell types (excitatory and inhibitory) distributed in four layers, and represents the cortical network below a surface of 1 mm^2^. The original implementation of the Potjans-Diesmann model was based on the NEST simulator and our goal here was to re-implement the model in the Brian 2 simulator and obtain the same results presented in the reference article. We did not replicate analyses that involve changes in the network structure. Our replicated network model presents activity dynamic patterns very similar to the ones observed in the original model, with comparisons made in terms of firing rates and synchrony and irregularity measures. In conclusion, the Potjans-Diesmann model was successfully replicated in a different platform than the one in which it was originally implemented.

## 2 Introduction

Most theoretical studies of cortical activity are based on networks of randomly connected units [2,6,7,12] or with architectures artificially built from random networks [10]. In spite of the usefulness of these models, in order to understand the interplay between network structure and cortical dynamics it is essential to have computational models which accurately represent the cortical network architecture. Recently, Potjans and Diesmann [8] developed a network model of the local cortical microcircuit based on extensive experimental data on the intrinsic circuitry of striate cortex [1,9]. The model contains two cell types (excitatory and inhibitory) distributed over four layers, L2/3, L4, L5, and L6, and represents the cortical network below a surface area of 1 mm^2^ (a scheme is shown in Fig. 1).

**Figure 1.**
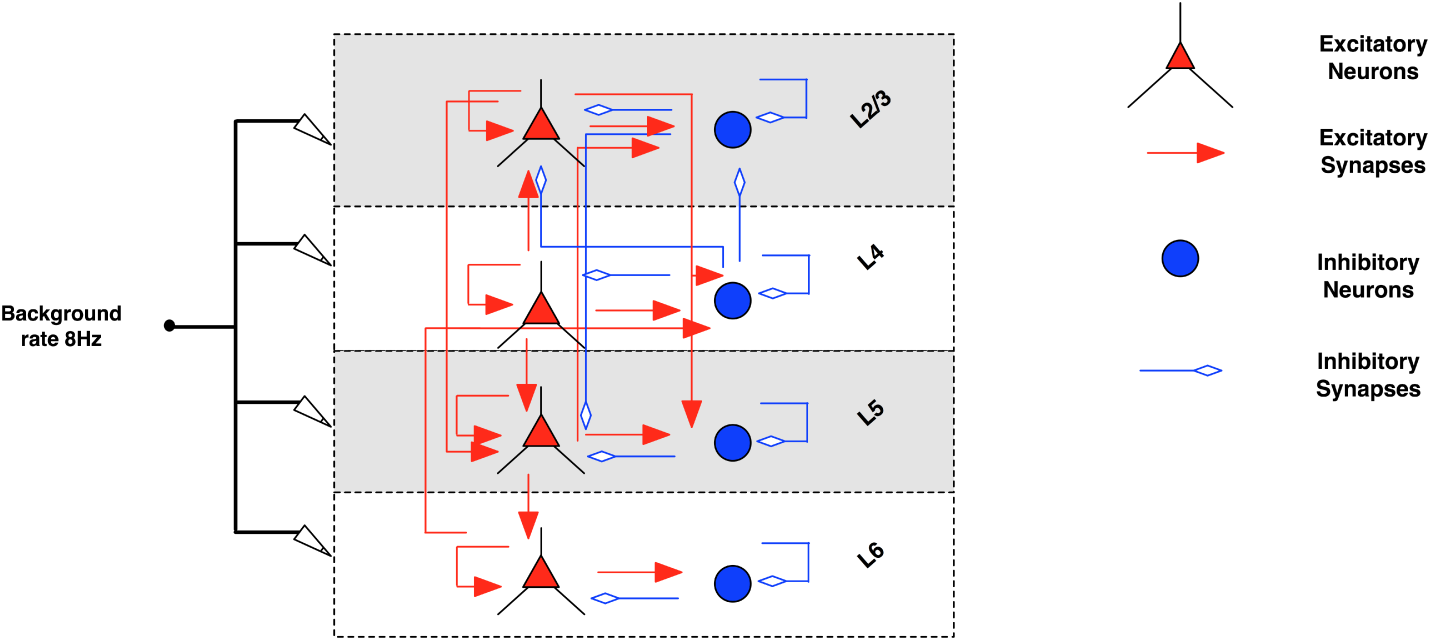
Schematic representation of the cortical network model (adapted from [8]). The model consists of four layers (L2/3, L4, L5 and L6), each one populated with excitatory (triangles) and inhibitory (circles) neurons (Table 2). Arrows represent connections with probabilities > 0.04: excitatory in red and inhibitory in blue (Table 3). Black arrows represent background inputs.

The original implementation was based on the NEST simulator [4] and the source code is available at the Open Source Brain platform [11]. Here, we reimplemented the full model in the Brian 2 simulator [5] without direct reference to the original source code.

## 3 Methods

In this work, we replicated in Brian 2 every detail of the Potjans-Diesmann model as described in their original article [8]. Hereafter, we will refer to the original NEST implementation of the Potjans-Diesmann model [8] as reference (or original) article. In this section we explain how this reimplementation was done. Further statistical analyses were performed using SciPy, NumPy, and Matplotlib libraries for the Python language.

### 3.1 Neurons

Network neurons are described by the leaky integrate-and-fire (LIF) neuron model. The subthreshold membrane voltage of neuron *i* obeys the equation

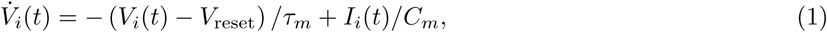

where *τ*_*m*_ is the membrane time constant, *C*_*m*_ is the membrane capacitance, *V*_reset_ is the reset potential, and *I*_*i*_(*t*) is the total input current. When *V*_*i*_(*t*) ≥ *V*_*th*_ the neuron emits a spike and the voltage is reset to *V*_reset_, remaining fixed at *V*_reset_ for a refractory period *τ*_ref_. The total input current *I*_*i*_(*t*) is divided into external *I*_*i*, ext_(*t*) and synaptic *I*_*i*, syn_(*t*). Whenever an excitatory (or inhibitory) neuron *j* presynaptic to neuron *i* fires at time 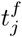, the synaptic current to neuron *i* changes at time 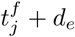 (or *d*_*i*_) by an amount *w* (or –*gw*), where *w* is the excitatory synaptic weight, *g* is the relative weight of the inhibitory synapse, *d*_*e*_ is the excitatory synapse transmission delay, and *d*_*i*_ is the inhibitory synapse transmission delay. In the absence of synaptic inputs the synaptic current changes as

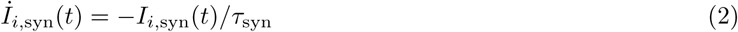

where *τ*_syn_ is the postsynaptic current time constant (parameter values are shown in Table 1).

**Table 1.** Neuron and synaptic parameters. Neuron and synaptic parameters used in our simulations according to [8].

|  |  |  |
| --- | --- | --- |
| membrane time constant | $\tau_m$ | 10 ms |
| refractory period | $\tau_{\text{ref}}$ | 2 ms |
| postsynaptic current time constant | $\tau_{\text{syn}}$ | 0.5 ms |
| membrane capacitance | $C_m$ | 250 pF |
| reset voltage | $V_{\text{reset}}$ | -65 mV |
| threshold voltage | $V_{\text{th}}$ | -50 mV |
| excitatory synaptic weight | $w$ | $\mathcal{N}(\mu = 87.8, \sigma = 8.8)$ pA |
| relative inhibitory synaptic weight | $g$ | 4 |
| excitatory synaptic transmission delay | $d_e$ | $\mathcal{N}(\mu = 1.5, \sigma = 0.75)$ ms |
| inhibitory synaptic transmission delay | $d_i$ | $\mathcal{N}(\mu = 0.8, \sigma = 0.4)$ ms |
| initial membrane potential | $V_0$ | $\mathcal{N}(\mu = -65.0, \sigma = 6.5)$ mV |

### 3.2 Network

The procedure to set up the network connections is the following:

- Start with a set of neurons *N* = 77,169, with model parameter described in Table 1.
- The *N* neurons are distributed over the eight different populations, L23e, L23i, etc, according to the numbers shown in Table 2.
- For each one of the sixty-four possible combinations of two from the eight cell populations, the total number *K* of synapses is calculated using equation (3) (compare with equation (1) of the original article), where *N*_*pre/post*_ are the sizes of presynaptic/postsynaptic populations and *C*_*a*_ is the corresponding connection probability given in Table 3 (the subindex *a* stands for ‘anatomical’ in the terminology of the original article).

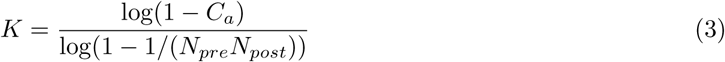
- For every one of the sixty-four two-cell populations, the *K* synapses determined above are created by uniformly and randomly choosing *K* pairs of neurons (one from each population) and placing a connection between them. This is done with repetition to allow the creation of multiple synaptic contacts between any pair of neurons.
- The synaptic weight for connections originating from excitatory neurons is set to w and the synaptic weight for connections from inhibitory neurons is set to –*gw*. In addition, the synaptic weight for connections from neurons of layer L4e to L23e is doubled [8,13].

**Table 2.** Layer population sizes (extracted from [8]). Neurons were distributed over four different layers (L23, L4, L5 and L6), and for each layer they were divided into excitatory (L23e, L4e, L5e and L6e) and inhibitory (L23i, L4i, L5i and L6i) subpopulations.

| L23e | L23i | L4e | L4i | L5e | L5i | L6e | L6i |
| --- | --- | --- | --- | --- | --- | --- | --- |
| 20683 | 5834 | 21915 | 5479 | 4850 | 1065 | 14395 | 2948 |

**Table 3.**
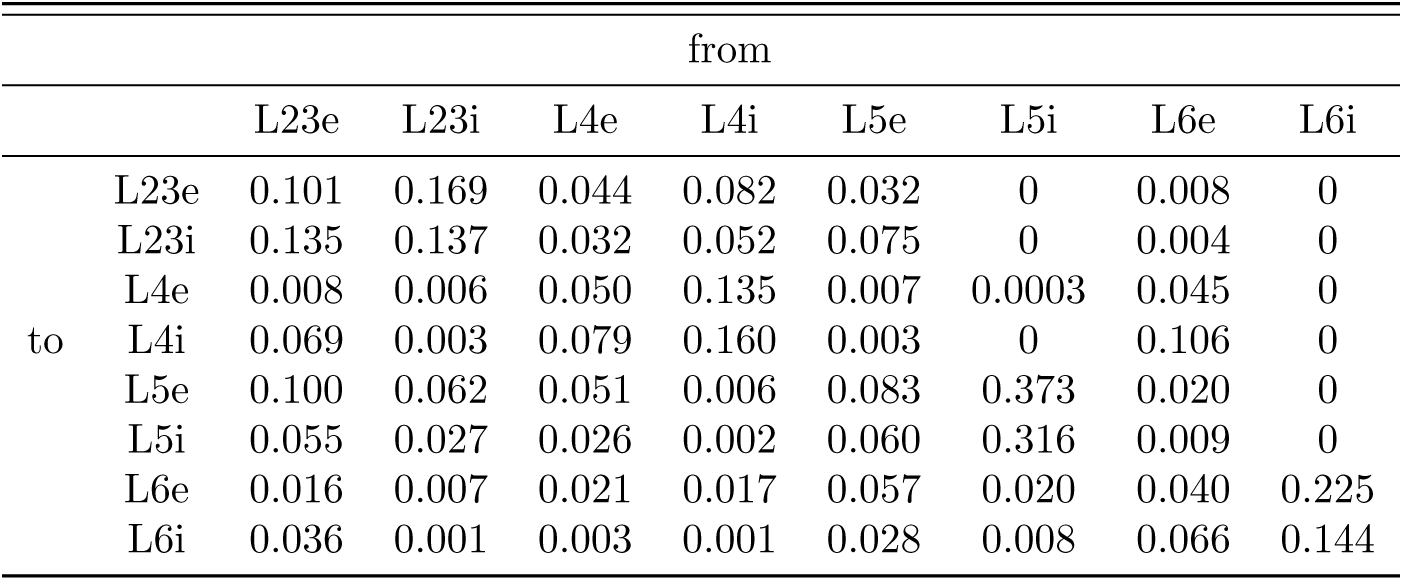
Connectivity matrix between the different populations of the model (extracted from [8]). The connectivity matrix describes the probabilities of the target-specific connections between populations of neurons.

The ordinary differential equations were solved with the exact integration method for linear equations available in Brian 2 with a time step Δ*t* = 0.1 ms. All simulations were carried out in a cluster with 4 nodes each equipped with 2 Intel Xeon processors.

### 3.3 External input

We chose from the reference paper three different types of external (“background”) inputs:

1. **Layer specific**: Neurons from each layer receive specific background spike-trains drawn from an 8 Hz Poisson distribution. The number of inputs per neuron is given in the first row of Table 4.
2. **Layer independent**: Spike-trains are drawn from an 8 Hz Poisson distribution as above, but now the number of inputs per neuron is the same for all the excitatory layers and the same for all the inhibitory layers as shown in the second row of Table 4.
3. **DC input**: The Poissonian background is replaced by constant DC currents to all neurons. The number of inputs per neuron follows the layer specific configuration (first row in Table 4). Observe that the number of inputs per neuron is multiplied by an effective factor which is given by *v* × *ω* × *τ*_*syn*_ = 0.3512 pA.

**Table 4.** Estimated numbers of external inputs per neurons in all network layers. The total number of external inputs is rounded.

|  | L23e | L23i | L4e | L4i | L5e | L5i | L6e | L6i |
| --- | --- | --- | --- | --- | --- | --- | --- | --- |
| Layer specific | 1600 | 1500 | 2100 | 1900 | 2000 | 1900 | 2900 | 2100 |
| Layer independent | 2000 | 1850 | 2000 | 1850 | 2000 | 1850 | 2000 | 1850 |
| Background rate ($\nu$) | 8 Hz | | | | | | | |

Additionally to these inputs, in Fig. 5C we simulate the network in several trials where the number of external inputs change. In every trial, the external inputs to each of the excitatory layers is a number randomly drawn between the layer specific and the layer independent inputs reported in Table 4. The inhibitory layers have their number of external inputs randomly decided between the already chosen number of their respective excitatory layer and the number calculated by

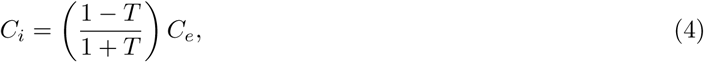

where *T* is the target specificity defined in [8] and is assumed here to be *T* = 0.1. However, there is an exception to this rule: L6i is allowed to have *T* = 0.2 due to the high number of inputs to L6e.

### 3.4 Measures

Here, we define the measures used to characterize the layer-specific activity of the network. They are the same ones used in the original article.

The spike train of a neuron *i* is represented by a sequence of temporal events (sum of delta functions). The firing rate over an interval *T* is obtained by summing the number of spikes during that interval and dividing by *T* (spike-count firing rate). The average firing rate of a population of *N* neurons is computed by calculating the firing rates of the neurons and dividing by *N*. With this procedure we calculated the average firing rates of the eight populations in the network.

To characterize irregularity in the network we use the coefficient of variation (CV) of the interspike interval (ISI) distribution. The CV_i_ for each neuron *i* is computed as the ratio of the standard deviation over the mean of its ISI distribution. Exponential distributions have CV ≈ 1 while more regular distributions have CV < 1.

Synchrony is characterized by the variability of the histogram of population spiking activity (bin size = 3 ms). The synchrony index is computed as the variance of the spike count histogram divided by its mean.

The degree of asynchronous and irregular activity in a population is quantified by a measure called AIness%. This is the percentage of the population with mean firing rate < 30 Hz, irregularity between 0.7 and 1.2, and synchrony < 8.

To compare the distributions of firing rates and CVs obtained from simulations in Brian 2 and NEST we use the Kolmogorov-Smirnov statistical test. To apply this non-parametric test, cumulated histograms were constructed with bins chosen by the Doane method [3].

## 4 Results

In the following, we present results of the replicated studies (with the same data sampling sizes) done in the reference article for the network with parameters as defined in Methods. We did not replicate analyses which involve changes in the network structure.

### 4.1 Spontaneous Activity

The simulated spontaneous activity is asynchronous and irregular (Fig. 2**A**) and the cell-type specific firing rates are in agreement with the ones observed in the reference article, including the lowest rates for the excitatory cells of layers 2/3 and 6 and the highest rates for L5 cells (Fig. 2**B**). For all layers the inhibitory cell firing rates exceed the ones of excitatory cells. The firing rate variabilities (Fig. 2**B**) and the single-cell firing rate irregularities (Fig. 2**C**) are also similar to the ones reported in the original article. The irregularity measure is > 0.80 for all cell populations (Fig. 2**C**). The profile of the synchrony of spiking activity across the cell populations is also consistent with the one reported in the reference article. The highest degree of synchrony is found in L5e and the lowest one in L6.

**Figure 2.**
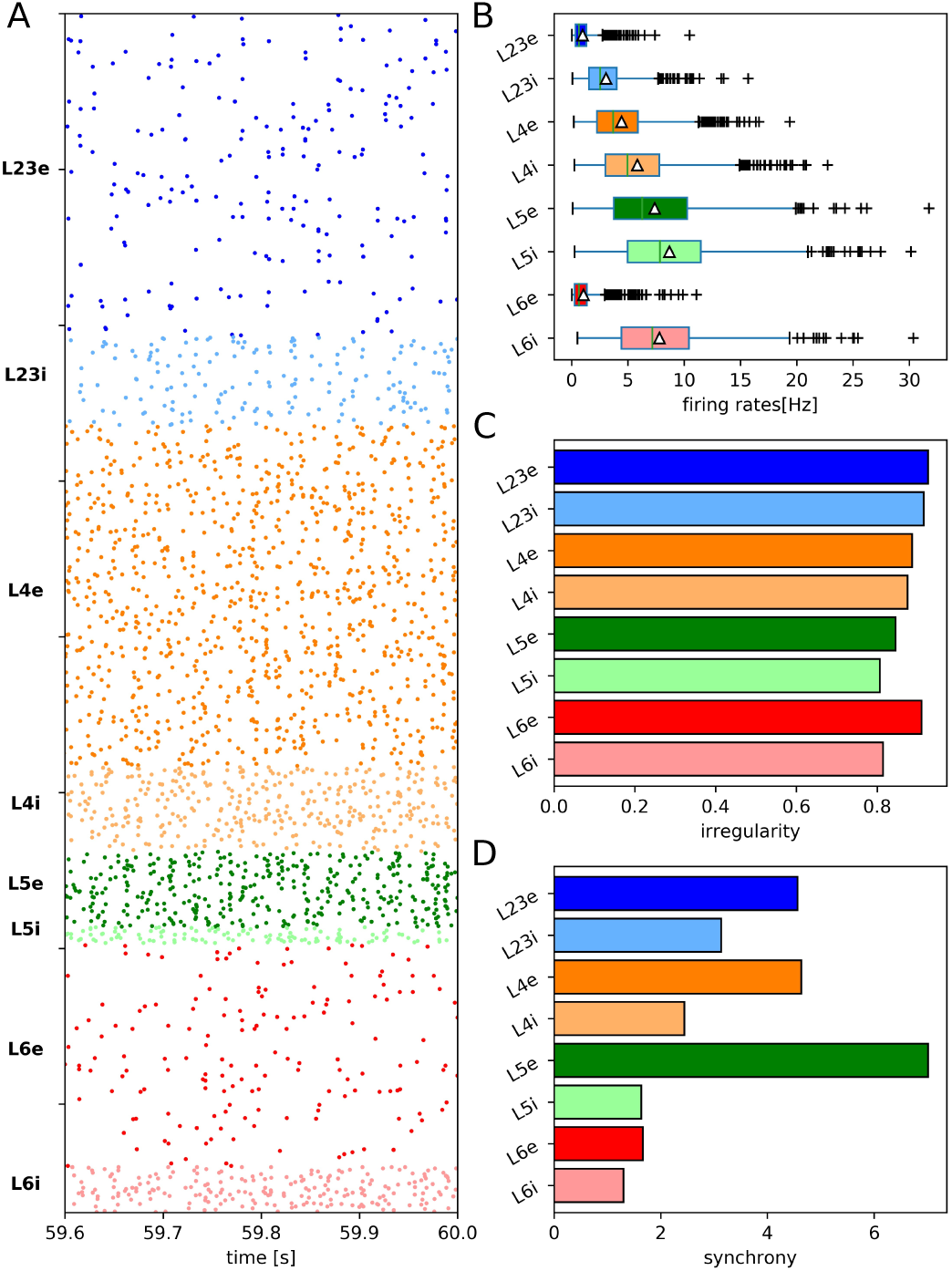
Spontaneous cell-type specific activity (to be compared with Fig.6 from the original article). In this simulation, *g* = 4 and background rate is layer-specific (8 Hz). (A) Raster plot of spiking activity for 0.4 s of all cell layers (from top to bottom; dark color: Excitatory cells, light color: Inhibitory cells). Number of displayed spike trains corresponds to 2.5% of the total number of neurons (preserving relative number of cells per layer). (B-D) Statistics based on samples of 1000 neurons per layer recorded for 60 s. (B) Single-cell firing rate boxplot. Triangles indicate population mean firing rates, crosses show outliers. (C) Irregularity of single-cell firing rates. (D) Synchrony of spiking activity.

In order to compare the Brian 2 with the NEST implementations of the Potjans-Diesmann model, we used the available pyNEST code [11] to run NEST simulations of the model with the same parameters given in Methods. The comparisons were made in terms of the cumulative distributions of the CVs of ISIs and the firing rates of the eight cell populations in the network (plots not shown here but as in Figs. 3 and 4 below). The p-value (two-sample Kolmogorov-Smirnov test) was higher than 0.6 for all comparisons meaning that no significant difference was found between NEST and Brian 2 simulations.

**Figure 3.**
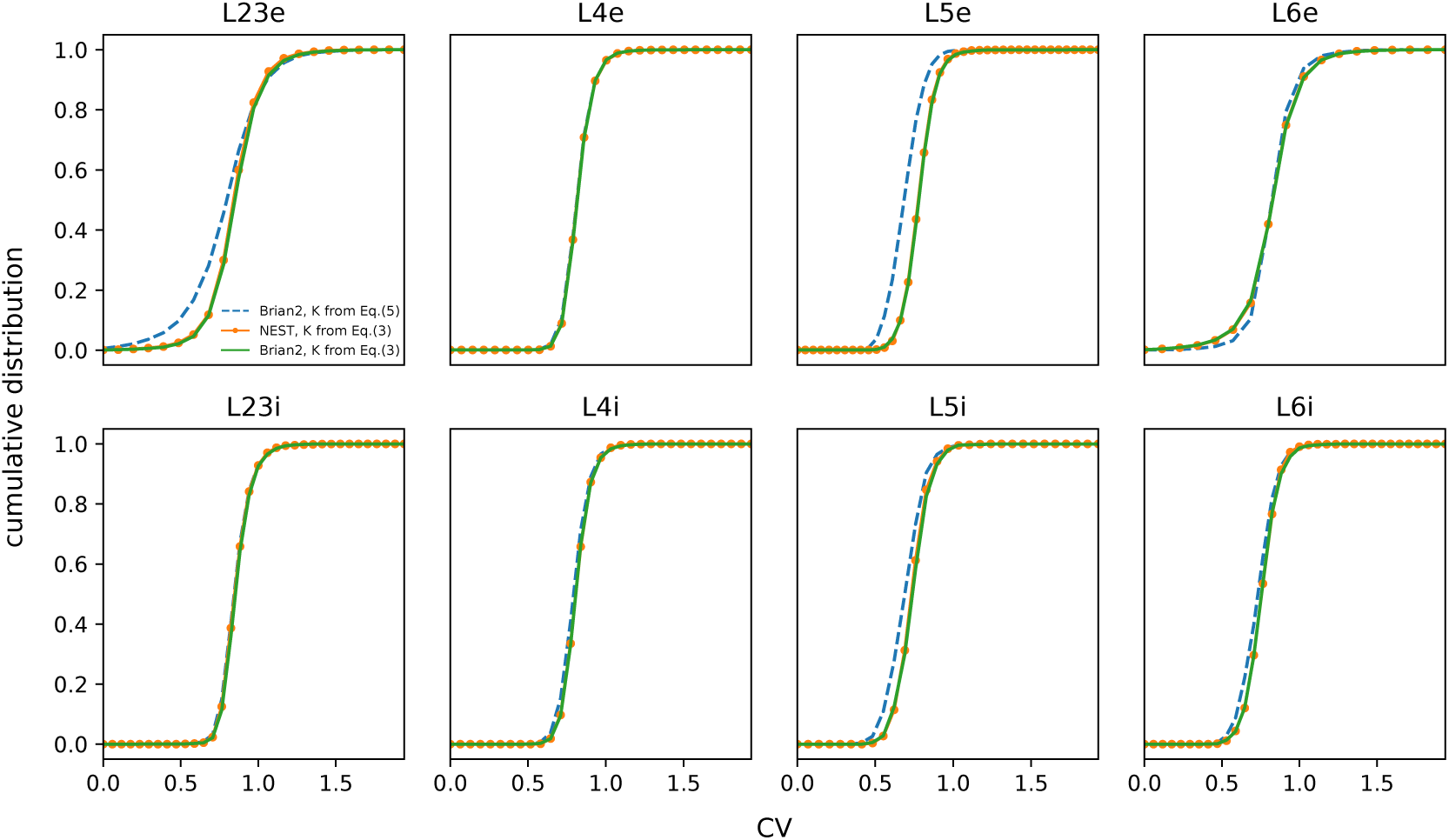
Cumulative distribution of CVs of ISIs for the eight cell populations (indicated atop each panel) for simulations implemented in Brian 2 using equation (5) (blue), in Brian 2 using equation (3) (green), and in NEST using equation (3) (orange). NEST code taken from [11].

**Figure 4.**
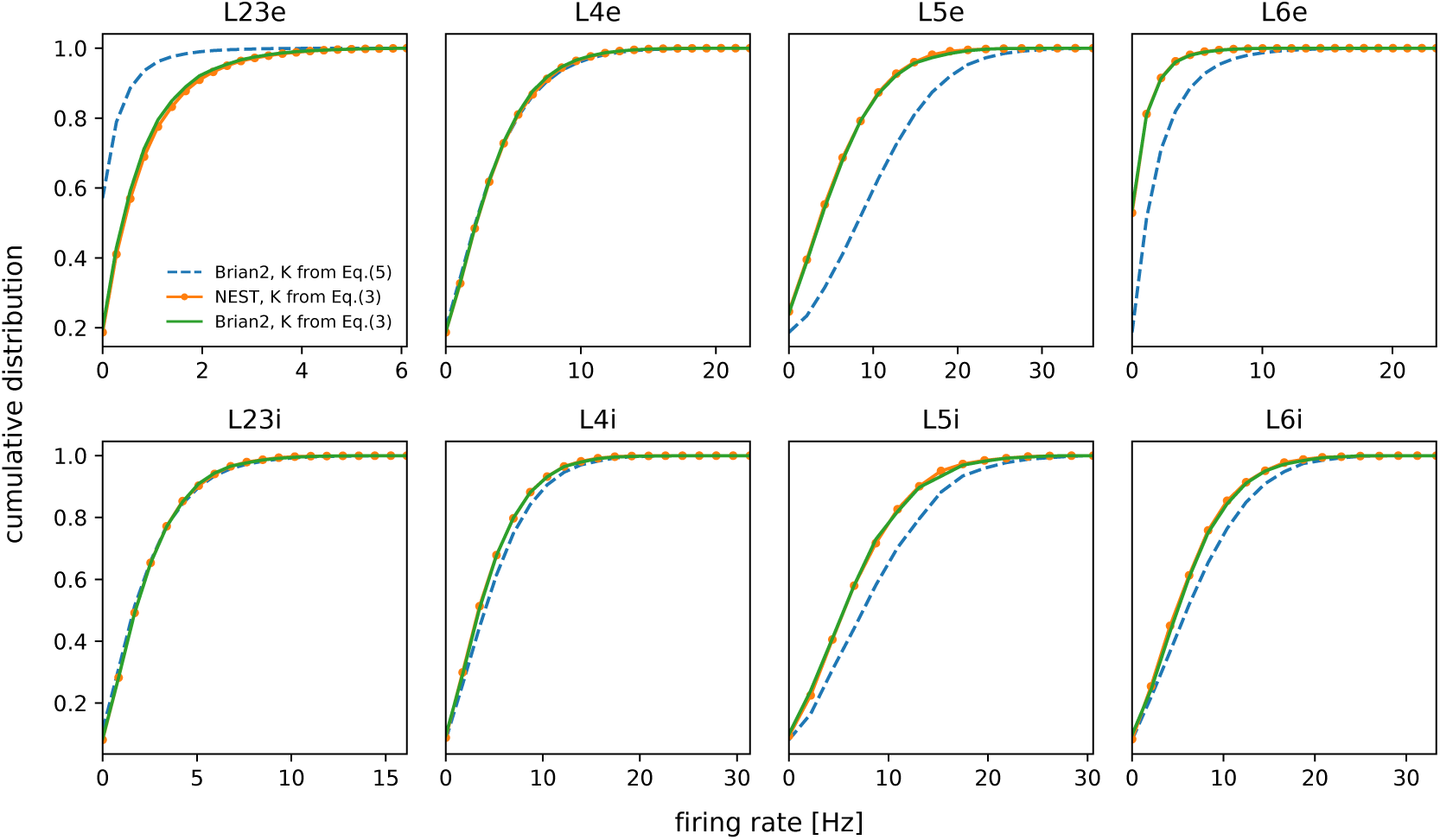
Cumulative distribution of firing rate for the eight cell populations (indicated atop each panel) for simulations implemented in Brian 2 using equation (5) (blue), in Brian 2 using equation (3) (green), and in NEST using equation (3) (orange). NEST code taken from [11].

Besides the creation of the network connections by the procedure described above, which is the one used in the original article, it is also possible to connect the neurons in the network using the alternative expression for the total number of synapses between two cell populations,

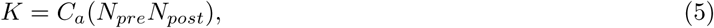

which comes from the first-order Taylor series approximation to the connection probability *C*_*a*_ as explained in the original article (see equation (2) and following text in the original article).

To test for possible effects of using this approximate equation, we constructed the network in Brian 2 using both equations (5) and (3). We found that the use of the approximate equation brings discrepancies in comparison with the original model. The comparisons were made in terms of the cumulative distributions of the CVs of ISIs (Fig. 3) and the firing rates (Fig. 4) of the eight cell populations for simulations of the original model based on the pyNEST code [11] and the Brian 2 code with K calculated from equations (5) and (3).

When comparing the network constructed in Brian 2 with equation (5) and the one constructed in NEST, most of the layer-specific comparisons yielded p-values higher than 0.6 as before, indicating no significant differences. However, the p-value of the comparison between the CV of ISIs for the layer 5 excitatory neurons was 0.07, and the p-values of the comparisons between the firing rates of excitatory cells from layers 2/3 and 6 were, respectively, 0.10 and 0.08. Therefore, in the cases of excitatory cells of layers 2/3, 5 and 6 there were statistically significant differences in the firing properties of neurons. The major discrepancy was found for L5e cells, which in the replicated Brian 2 version using equation (5) have firing rate 12.2 ± 6.1 Hz while in the original version their mean firing rate is 7.8 ± 5.1 Hz.

The differences found highlight the importance of using the exact expression for the connection probability *C*_*a*_ given in equation (1) of the original article (which corresponds to equation (3) of this text) in simulations of the Potjans-Diesmann model. Observe the close agreement of the cumulative histograms when Brian 2 (K from equation (3)) and NEST (K from equation (3)) are compared.

We now return to the reimplementation of the Potjans-Diesmann model in Brian 2 using equation (3), which will be kept for the rest of this replication work.

### 4.2 Dependence of Spontaneous Activity on External Inputs

In agreement with the results obtained in the reference article, the activity features of the Brian 2 implementation are also robust to changes in the external inputs. These are shown in Fig. 5A, in which the Poissonian inputs are replaced by constant DC currents, and in Fig. 5B, in which the layer-dependent Poissonian inputs are replaced by layer-independent inputs. In the latter case, the absence of activity in L6e resulting from the layer-independent inputs indicates the importance of realistic input structure to yield plausible activity in all layers. In Fig. 5C we present the population firing rates for 100 trials with the rule explained in the methods section. This latter experiment presented an excellent agreement with the histograms observed in Fig. 7 of the original article.

**Figure 5.**
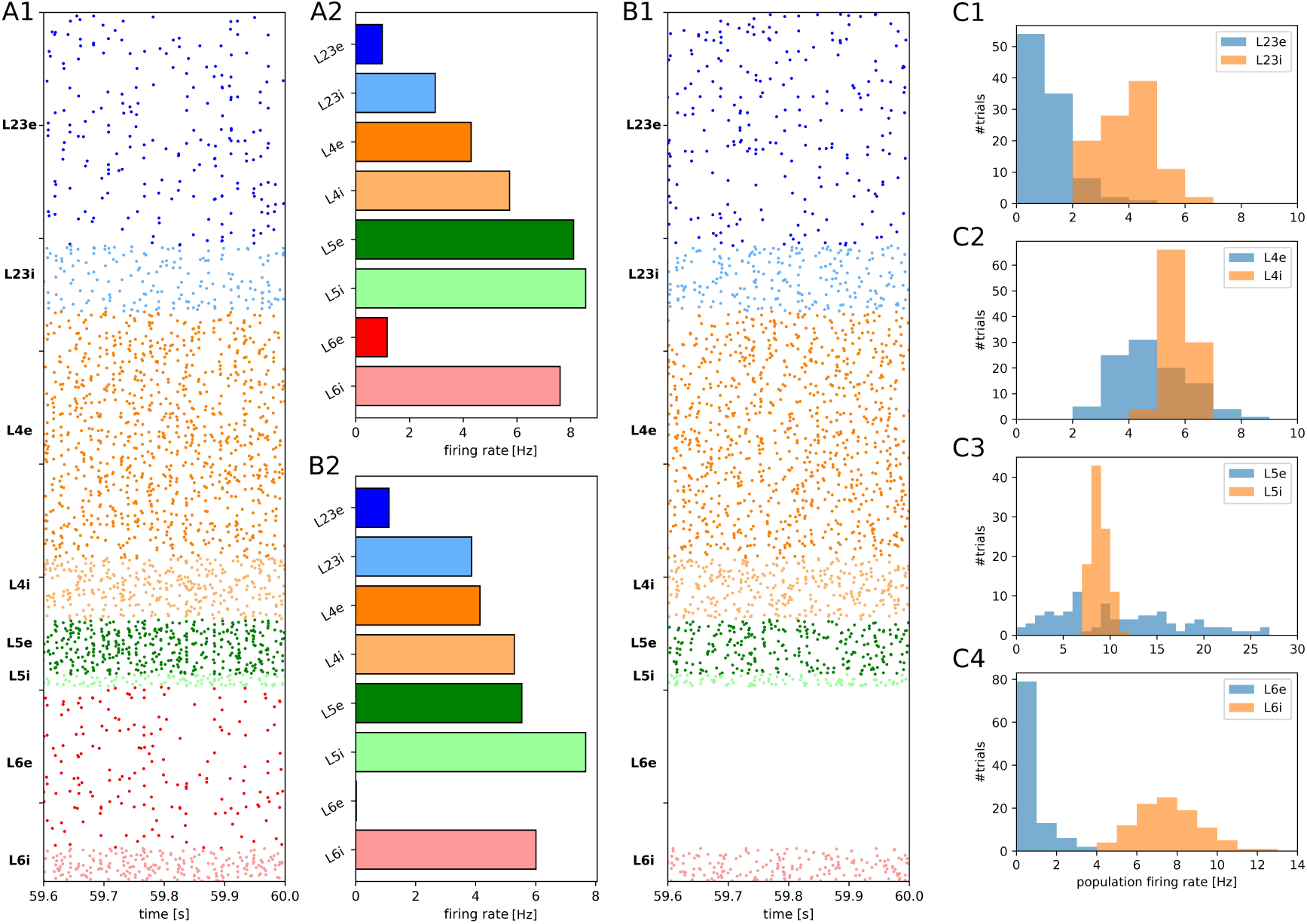
Raster plot of spiking activity over a 0.4 s period and average firing rate for different types of external input (to be compared with Fig. 7 of the original article). Color codes and sample cell sizes as in Fig. 2. (A1–A2) DC input. (B1–B2) Layer independent input. (C1–C4) Histograms containing the population firing rates of 100 trials where the external inputs were drawn with a specific rule (see details in methods).

### 4.3 Stability of Network Activity

The activity features obtained in the original article by changing the relative strength of inhibitory synapses and the background rate are reproduced in our reimplementation (Fig. 6).

**Figure 6.**
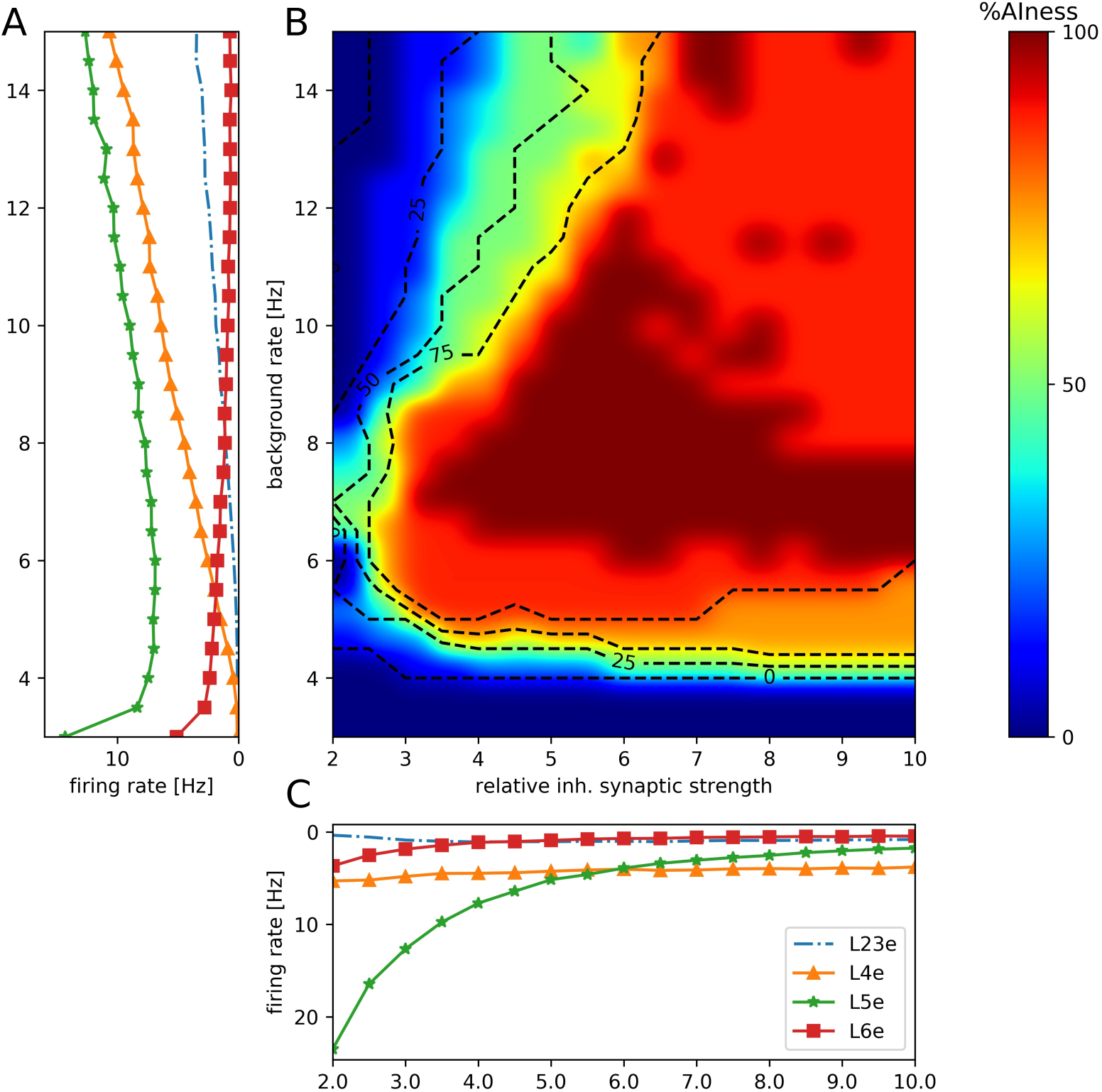
Network activity dependence on background rate and relative inhibitory weight (to be compared with Fig. 8 of the original article). (A) Mean population firing rates of excitatory neurons in layers 2/3 (dashed line), 4 (triangles), 5 (stars) and 6 (squares) as a function of the background rate for fixed *g* = 4. (B) AIness% (see Methods) as a function of the background rate and the relative inhibitory synaptic strength *g*. Labeled dashed contour lines indicate areas where 25%, 50% and 75% of all populations fire in an asynchronous and irregular mode at low rate. (C) Mean population firing rates of excitatory neurons as a function of the relative inhibitory synaptic strength g for fixed background rate 8 Hz (markers as in (A)).

The asynchronous and irregular activity of the reimplemented network model, as characterized by the AIness%, is similar to the one found in the reference article for background rates ¿5 Hz and relative inhibitory synaptic strengths ¿4 Hz. In comparison with the reference article, the relative order of excitatory firing rates is maintained for every combination between background rate and relative synaptic strength, with highest values in L5 and smallest in L2/3 and L6. Similarly to the original article, the firing rate of L4e cells is the most sensitive to variations in the background rate whereas the firing rate of L5e cells is the most sensitive to variations in the relative inhibitory synaptic strength *g*.

## 5 Conclusion

Using the Brian 2 reimplementation of the Potjans-Diesmann model we were able to reproduce the main results of the original article [8]. The spontaneous activity of the network reimplementation is asynchronous and irregular as evaluated by the different measures used to characterize spiking behavior.

We also have shown the importance of using the exact expression in equation (1) of the original article instead of the approximate one in equation (2) in implementations of the model. The use of the approximate expression leads to mean firing rates of L5e neurons significantly higher than in the original implementation of the model.

The successful replication of the results of the reference article confirms the correctness of the original implementation of the model.

## 6 Acknowledgements

The authors would like to thank Prof. Markus Diesmann for valuable comments on the manuscript. This research was supported by the São Paulo State Research Foundation (FAPESP) grant No.2013/07699-0 for the Research, Innovation and Dissemination Center for Neuromathematics, FAPESP grant 2016/03855-5 to NLK, FAPESP scholarship 2017/07688-9 to RS, FAPESP scholarship 2013/25667-8 to RP and FAPESP scholarship 2017/05874-0 to VC. ACR is partially supported by the CNPq fellowship grant 306251/2014-0. RP and ACR are also part of the IRTG 1740 / TRP 2015/50122-0, funded by DFG / FAPESP. CCC, and CR are supported by a CAPES PhD scholarship.

